# Using movement data to estimate contact rates in a simulated environmentally-transmitted disease system

**DOI:** 10.1101/261198

**Authors:** Eric R. Dougherty, Dana P. Seidel, Colin J. Carlson, Wayne M. Getz

**Affiliations:** Department of Environmental Science Policy and Management, University of California, Berkeley; Schools of Mathematical Sciences, University of KwaZulu-Natal, South Africa

**Keywords:** Agent-based Model, Infectious Disease, Environmental Transmission, Contact Rate, Animal Movement, Anthrax

## Abstract

Agent-based models have become important tools in ecology, particularly in the study of infectious disease dynamics. Simulations of near-continuous movement paths guided by empirical data offer new avenues of investigation into disease transmission. Here, we simulate the spatiotemporal transmission dynamics of anthrax, the acute disease caused by the bacterium *Bacillus anthracis*, a pathogen transmitted primarily via environmental reservoirs. We explore how calculations of the probabilities of contact between a host and infectious reservoirs are affected by the scale and method of analysis. At both the landscape and individual scales, empirical movement tracks offer previously unattainable estimates of impacts of movement decisions on contact rate metrics. However, the analytical method selected for the calculation of the probability of contact has notable impacts on the resulting estimates, with convex polygons virtually canceling out variation, and unions of local convex hulls (LoCoH methods) and space-time prisms reflecting reasonable variation, but differing in the magnitude of their estimates. The explicit consideration of behavioral states along movement pathways also impacts evaluations of exposure risk, though its effects differ across methods of analysis. Ultimately, simulations demonstrate that the incorporation of movement data into pathogen transmission analyses helps clarify the role of movement processes underlying the observed dynamics of infectious disease.

## Introduction

Though epidemic dynamics are often simplified using rates averaged at the population scale [1], the transmission process underlying disease spread is highly dependent on various forms of heterogeneity [2]. The influence of such heterogeneity has been recognized in both human [3] and wildlife [4] populations, but the diversity of underlying causes of individual variance often makes precise measurement of contributing factors difficult. The impact of any one individual on the subsequent dynamics of an epidemic arises from a complex combination of host and pathogen characteristics, as well as the environment in which transmission occurs [3]. A variety of methods, often applied during or after an epidemic, have demonstrated this variation in natural systems [5, 6]. However, predicting *a priori* which forms of heterogeneity will result in individuals who contribute disproportionately more secondary infections than the average infected individual (i.e., superspreaders; see [3] for details) represents a notable challenge in disease ecology [7].

One fundamental driver of heterogeneity in disease transmission is the variance in movement behaviors among host individuals [8, 9]. Until recently, however, the tools available to observe empirical movement trajectories did not offer fine enough resolution to describe individual heterogeneity beyond differences in space-use at landscape scales. With advancements and innovations in bio-logging technologies, researchers have gained the ability to identify specific behavioral states in movement tracks and resolve general distributions that give rise to heterogeneity among individuals [10]. Even so, the consideration of heterogeneous movements remains relatively underexplored as a major contributor to the unique dynamics that characterize epidemic expansion or fade-out (i.e., when the introduction of a disease fails to propagate widely) [9]. As movement data becomes more readily available in disease research and the methods for analyzing such data continue to develop, the incorporation of this form of heterogeneity in disease models will be vital for accurately reflecting spatiotemporal disease spread.

Agent-based models (ABMs) represent a direct means of incorporating individual heterogeneity into disease models [11]. ABMs refer to computational simulation models consisting of interacting components (i.e., agents and their environment) that follow a set of explicit rules. These general rules enable the agents to move independently as they seek to fulfill specific objectives and can respond to various stimuli in their surroundings, whether this comes in the form of environmental changes or interactions with other agents. Rather than imposing certain expectations on the output of the model, this bottom-up approach allows for the natural emergence of patterns at broader levels of analysis [12, 13]. The flexibility of this framework has led to broad applications of ABMs even within the ecological literature, including to questions in environmental resource management [14, 15], examinations of evolutionary dynamics [16, 17, 18, 19], and considerations of individual animal behaviors [20, 21]. Unlike compartmental models (the traditional means of modeling epidemic dynamics), which assume homogeneous mixing within compartments, ABMs offer a framework in which heterogeneity can emerge as it does in natural systems.

ABMs have also increasingly been used to explore disease dynamics, with individual agents often transitioning between the infectious stages normally indicative of a compartmental SIR model (e.g., susceptible, infectious, and recovered [22]). The general rule set of an ABM typically gives rise to stochasticity, and thus heterogeneity, in behaviors such as movement. This makes ABMs ideal tools for exploring the role of individual movement decisions in the propagation of disease as a stochastic process. In the few cases that have utilized ABMs to explore disease systems, individual movement dynamics are often simplified to diffusion processes [23, 24] or highly generalized jumps between patches [25, 26, 27]. Though other models have implemented more complex mechanistic movement rules to govern agent trajectories (e.g., [28, 29, 30, 31, 32]), examples of such ABMs remain fairly limited in disease ecology, and their applicability is frequently constrained by the specific nature of pathogen transmission in the focal disease system.

Here we develop an ABM for herbivore movement on an anthrax-endemic landscape, parameterized with empirical movement data. As anthrax is transmitted primarily through environmental reservoirs [33, 34, 35], we use the output of the model to estimate the probability of contact between agents and fixed environmental reservoirs (anthrax spores deposited at carcass sites), an important epidemiological rate that is highly dependent on the heterogeneity in individual movement. Rather than explicitly simulating the disease transmission process, we only aim to estimate contact rates between a host and an infectious agent. Doing so allows us to illustrate (and critically examine) the utility of movement ecology in disease research, without having to correct for the complex disease dynamics that would emerge if we included transmission in the model. High-resolution GPS data offers insight into the general rules that dictate host movements, enabling an accurate consideration of the heterogeneity in this fundamental contributor to the disease transmission process. Furthermore, applying analytical tools from movement ecology offers the ability to evaluate variation in an important epidemiological process at the population level, despite collecting data on a relatively small subset of the population at risk. Thus, we demonstrate the multi-faceted manner by which the data and methods from movement ecology can contribute to our understanding of the heterogeneity that defines disease systems.

## Simulation Methods

Anthrax, the acute disease caused by the bacterium *Bacillus anthracis*, remains a persistent threat in many wildlife populations throughout the world [36, 37]. Though a variety of animal species can contract the zoonotic disease, herbivores experience the highest mortality rates, while many carnivores and scavengers exhibit resistance or tolerance [37, 38]. In some systems, anthrax outbreaks are seasonally-driven, though there may exist inter-specific differences in the timing of the peak of infections. For example, zebra in Etosha National Park in Namibia experience peaks in infection during the warm wet season (March-April), whereas elephants are more likely to be infected during the dry months of October-November [35]. A definitive explanation for these peaks remains elusive, but a number of alternatives have been proposed, including nutritional stress, heterogeneous soil ingestion rates [33], and complex coinfection dynamics [39, 40].

*B. anthracis* takes the form of reproducing vegetative cells in infected hosts and endospores when in soil and ponded water environments [36, 41], although some vegetative reproduction may take place within the rhizosphere of vigorously growing grasses [42]. The spores are exceptionally resilient in the face of environmental stress, and allow the infectious agent to persist in environmental reservoirs for extended periods of sub-optimal conditions [41]

Spores can enter the host organism through cutaneous lesions, by inhalation into the pulmonary system, or via the gastrointestinal (GI) tract. Many ungulates consume substantial amounts of soil in addition to vegetation during foraging bouts, and in doing so, may inadvertently ingest the pathogen [33]. Limited evidence from necropsies suggests that GI infections are the most common route of infection in herbivores [43] and will be the primary mechanism modeled here. Anthrax is highly pathogenic in herbivores, and death may occur within a few days or up to two weeks after contact with a lethal dose of *B. anthracis* spores [35].

Anthrax is endemic in the plains herbivores of Etosha National Park, Namibia, peaking in zebra, springbok, and wildebeest during the rainy season and in elephants during the dry season [33, 44, 40]. Extensive carcass surveillance efforts in Etosha National Park, Namibia, between 1968 and 2011, conducted by The Etosha Ecological Institute [33, 45, 38], were used to inform the densities of anthrax infected carcasses considered in the simulation. In addition, empirical movement data collected from Etosha National Park, were used to inform the agent components of the simulation model. Specifically, the movement trajectories of nine zebra (Table 1) were used to estimate 20-minute step-length and turning angle distributions, as well as the general activity budget (i.e., the distribution of GPS fixes falling into each of the three possible behavioral states based on all of the points obtained from zebra). We used both carcass and movement data to develop a simulation model and explore the role of individual movement in disease transmission. Our description follows the ODD (Overview, Design concepts, and Details) framework [46].

### Purpose

The simulation model consisted of agents (with size and movement characteristics obtained from zebra data) moving across a simulated heterogeneous landscape upon which infected carcasses (three sizes with biomasses corresponding to springbok, zebra, and elephants) were deposited at the initialization of the model. A probability of contact with locally infectious zones (LIZs; [47]) was calculated for each individual throughout a single anthrax season (defined as the three month period between February and April; [33]). This probability serves as a synthesized epidemiologically-relevant metric that is then estimated using alternative movement analyses.

### State Variables and Scale

The three fundamental interacting components of our agent-based model are: 1) the landscape, which reflects the forage quality across the simulated park; 2) the agents that represent susceptible ungulates moving according to empirically-driven rules; and 3) the locally infectious zones, which are the infectious component of the model and form the basis of the transmission dynamics. The state variables of interest for each of these components are displayed in Table 2 and are explained in more detail below.

A heterogeneous landscape of forage quality was established to serve as the mechanism driving the movement of agents. A square study area covering 2,500 km^2^ (50 km by 50 km) was created. Hexagonal cells, each with a radius of 200 meters, were placed atop the study area, defining the landscape under surveillance. The edges of the hexagonal grid act as a fence and prohibit movement outside of the study area. Though this imagined area is substantially smaller than the entire extent of Etosha National Park (approximately 22,000 km^2^), it is consistent in scale with the mean 100% minimum convex polygon (MCP) of the empirical movement tracks, which was 2522.8 km^2^. Though the maximum 100% MCP was larger than 2500 km^2^, the fact that all of the empirical tracks had durations longer than three months (the length of the anthrax season) suggested that the mean MCP area would serve as a reasonable extent for our simulation.

Agents, representing individual ungulates moving across the landscape, were assigned a body size that did not change over the course of the simulation (*B*_*i*_) and a perceptual range parameter (*P*_*i*_), according to the following equations:

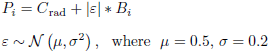

In this way, each individual perceives at least the cell in which it was currently located, but may have been able to perceive larger areas, with larger individuals likely having increased ability to perceive the quality of cells further away [48]. In addition, each agent was assigned to a particular behavioral state following the distribution of behavioral states observed in the nine empirical movement trajectories from Etosha National Park based on a Hidden Markov Model (HMM; [49]; see Supplementary Materials). The three possible states were resting (State 1), foraging (State 2), and directed movement (State 3). These states were associated with particular characteristics of movement, described below.

Locally infectious zones (LIZs), centered around the point locations at which an animal succumbs to its infection, are the critical infectious components of the anthrax system [35, 33, 34]. Due to the resilience of the *B. anthracis* spores, the area immediately surrounding an infected carcass can contain infectious material for extended periods of time (on the order of multiple years) [50]. Subsequent visits by grazers to these LIZs may result in their infection when spores in the soil are incidentally ingested along with vegetation [33]. These LIZs were identified in the model as either small (representing a springbok-sized carcass), medium (representing a zebra or wildebeest carcass), or large (representing the occasional elephant-sized carcass) according to the general distribution observed in empirical carcass data from Etosha National Park. Each LIZ was then assigned an initial mass (associated with the individual at the time of its death) based on the size category to which it was designated. In addition, the age of the LIZ (the number of years prior to the initialization of the model, up to three) was assigned, such that the number of the LIZs on the landscape deposited each year were approximately equal and reflected empirical observations in Etosha National Park (and considered with the error associated with surveillance efforts; [45]). The latter state variable was used to determine the relative increase in attractiveness associated with the landscape cell upon which the LIZ was deposited (see Initialization below).

### Process Overview and Scheduling

Following the initialization of the model, there are two primary sets of processes that occur during the execution of the simulation: the movement processes affecting the positions and behavioral states of agents and the landscape feeding (general and specific) and growth processes affecting the current forage quality values in various hexagonal cells. The scheduling of these processes is depicted in Figure 1.

**Figure 1:**
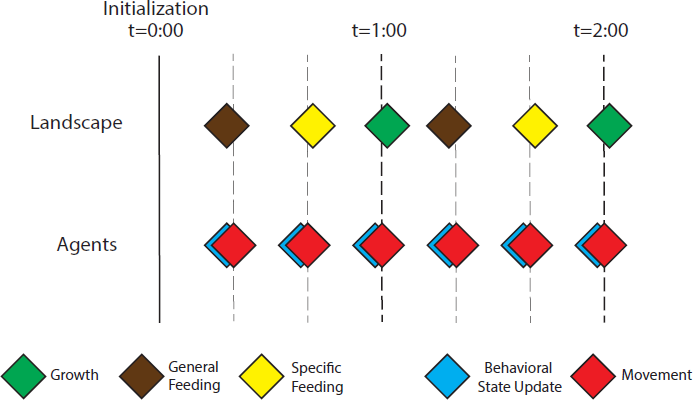
Process Diagram. The scheduling of the landscape and movement processes throughout the simulation. Following initialization of the landscape and agents (t = 0 : 00, hours:mins), the first process to occur is a general feeding bout whereby a set of *untagged individuals* (i.e., those with unknown movement pathways that are consequently randomly dispersed across the landscape) effectively decrease the current forage quality of underlying cells. Then the movement processes occur at the first 20 minute interval. First the behavioral state is evaluated and potentially shifted, then a step is taken based on the step size and turning angle distributions associated with the current behavioral state of each agent. The next behavioral state shift and positional change is undertaken prior to the next landscape process, specific feeding. This occurs immediately following the second move of the hour (i.e., at t = 0 : 40), and only cells with *tagged individuals* occupying them are affected. Another update to the behavioral state and position of the agents is executed before the final landscape process (growth) occurs, effectively beginning the same sequence of steps again for each hour of the 24-hour day over the course of the 90-day anthrax season.

### Design Concepts

**Emergence:** The key measure of the simulation, the individual probability of contact with infectious material, emerges from the mechanistic movements of the agents (during the foraging behavioral state) within the immediate vicinity of LIZs on the landscape. The probabilistic co-occurrence of the two components of the model give rise to the metric of interest.
**Sensing:** The perceptual range variable assigned to each agent governs the manner by which agents sense their surrounding. Agents are presumed to sense perfectly their own behavioral states, which dictates their subsequent movement.
**Interaction:** Interaction occurred only between agents and the landscape, and only during one of the behavioral states (foraging). An implicit interaction between LIZs and the landscape is also incorporated at initialization (in the form of their preferential placement on high quality patches), but this interaction is not continuously updated as in the case of the agent movements.
**Stochasticity:** Stochasticity was incorporated in multiple ways. The initial state variables were derived in a stochastic manner for all three components of the model (Landscape, Agents, and LIZs). In addition, the behavioral states of agents were stochastically generated using binomial draws with probabilities derived from empirically-driven transition functions (described below). Once the behavioral state was selected, the steps and turning angles were stochastic in that they were drawn from state-specific distributions.
**Observation:** The behavioral states of each individual over time were not observed directly, but they were theoretically extractable from the observed movement paths. Similarly, the actual forage quality layer, representing the mechanism underpinning agent movements in the foraging behavioral state, was not directly observable, and only a proxy akin to NDVI was available for downstream analyses (Figure 2).

**Table 1:** Serial location data (collected every 20 minutes from 9 zebra) that were used for parameterizing movement in the simulation model.

| Individual ID | Number of Points | Start Date | End Date |
| --- | --- | --- | --- |
| AG059 | 20,769 | 2009-04-20 | 2010-08-29 |
| AG061 | 17,196 | 2009-04-20 | 2010-03-03 |
| AG062 | 12,794 | 2009-04-20 | 2010-01-30 |
| AG063 | 25,721 | 2009-04-20 | 2010-04-30 |
| AG068 | 32,661 | 2009-04-20 | 2010-08-29 |
| AG252 | 23,450 | 2009-10-06 | 2010-08-29 |
| AG253 | 21,676 | 2009-10-06 | 2010-12-17 |
| AG255 | 23,470 | 2009-10-06 | 2010-08-29 |
| AG256 | 23,519 | 2009-10-06 | 2010-08-29 |

**Table 2:** Descriptions of the state variables associated with each of the components of the agent-based model.

| Component | State Variables | Description |
| --- | --- | --- |
| Landscape | Forage Carrying Capacity | The maximum carrying capacity for forage in each hexagonal cell |
|  | Current Forage Quality | The current value of the hexagonal cell following feeding and growth processes |
| Agents (zebra only) | Body Size | The size (in kilograms) of the individual agent; mean of 350, SD of 43.75 |
|  | Perceptual Range | The range (in meters) within which the agent perceives the quality of vegetation patches |
|  | Behavioral State | One of three possible movement states: resting, foraging, or directed movement |
| LIZs (springbok, zebra, wildebeest, elephant) | Species | One of three possible carcass size classes: small (springbok), medium (wildebeest, zebra), or large (elephant) |
|  | Initial Mass | The mass (in kilograms) of the carcass upon initial deposition based on the species classification |
|  | Time Since Deposition | The time (in years) since the animal had succumbed to anthrax and became a carcass-producing LIZ |

**Figure 2:**
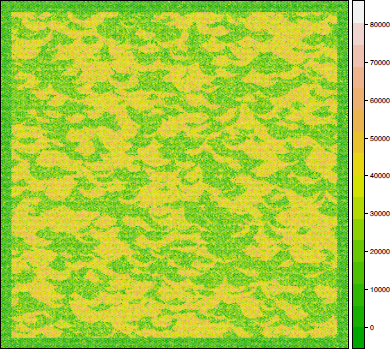
Initialized Landscape. A representation of the forage quality as perceived by the grazing agents on the landscape. An adjustment was made to decrease the quality of the forage near the fence along the outer boundary of the “park.”

### Initialization

Each landscape cell *j* = {1,…, 72250} was randomly assigned a relative forage quality value (*Q*_*j*_) that was dependent on an overall mean forage quality value for the landscape (*μ*_*F*_; in kg/km^2^). This value was set as a randomly selected value from a uniform distribution between 900,000 and 1,100,000 (based on estimates using satellite monitoring technology; [51]). Because each cell was considerably smaller than 1 km^2^, the overall mean forage quality that served as the basis for assigning a value to each cell was scaled accordingly (i.e., each cell contained an area of approximately 34640 m^2^, so the mean forage value was reduced to 0.03464 * *μ*_*F*_). Ultimately, the forage quality value of each standard cell (we also have high and low quality cells, as discussed below) emerged from the distribution *Q*_*j*_ ~ *N* (*μ*, *σ*^2^), where *μ* = *μ*_*F*_ * 0.03464 and *σ* = *μ*_*F*_ * 0.03464/4

Two percent of the cells were then selected as high quality cells and assigned a value from a normal distribution with a mean 1.33 times that used in a standard cell. A radius of influence was then assigned to each of the highest quality cells, equivalent to five times the hexagon cell size (*C*_rad_ ≈ 1000 meters) times an arbitrary factor chosen randomly from a normal distribution with a mean of 1.5 and a standard distribution of 0.5. The same process was carried out for selecting low quality cells, but the mean of the normal distribution was set as 0.66 that of a standard cell. The quality of each cell whose center fell within this radius of influence was then adjusted by selecting a value from the same high-quality normal distribution. Thus, each of the highest quality cells was surrounded by some variable number of cells with similar quality to effectively create patches of generally higher quality than the rest of the landscape (see Figure 2, left panel). These values represented the carrying capacity of that particular cell (*K*_*j*_), such that growth processes (described below) would return the quality of the cell to these levels when it was unoccupied by an agent. Following this, cells whose center fell within a buffer of 1500 meters of the edge of the park were reassigned a value drawn from a normal distribution with a mean of 0.33 times that of a standard cell.

At the beginning of the anthrax season (February 1), 18 agents were assigned an initial behavioral state using a random number drawn from a uniform distribution ranging between 0 and 1. The probability of beginning the simulation in State 1 was approximately 0.080 (i.e., a random value between 0 and 0.080), the probability of beginning in State 2 was approximately 0.542 (i.e., a random value between 0.080 and 0.622), and the probability of beginning in State 3 was approximately 0.378 (i.e., a random value between 0.622 and 1). All agents in behavioral state 2 were placed upon the landscape such that they were utilizing cells of high quality (i.e., cells with values greater than one standard deviation above the mean grazing value), but were placed randomly otherwise.

The placement of LIZs *k* = {1, …,*z*} was also dependent on the underlying forage quality layer. A slightly higher proportion (70%, rather than approximately 54%) of LIZs were placed on cells with forage quality greater than one standard deviation from the mean forage quality to reflect the fact that LIZs actively increase the quality of the cell in which they are deposited [34]. In addition, a “green up” was induced at initialization to increase the carrying capacity of cells containing anthrax carcasses (*K*_*j,t*_; where the index *t* was used set up an iterative relationship over time), thereby impacting the growth curve in the cell (see Landscape Processes, below). The amount of increase depended upon the age of the LIZ (*A*_*k*_), and followed the equation:

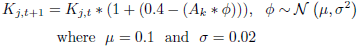

The initial mass of the carcass (*B*_*k*_) and the time since deposition (*A*_*k*_) were used to create a buffer around the central point associated with the LIZ. The size of the buffer (*S*) of each LIZ (*k*) was determined by the following equation:

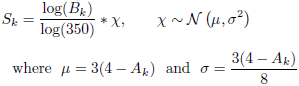

These polygons served as the synthesized LIZ layer for the calculation of the number of contacts (described below in Analysis Methods).

### Input

The input data used to inform the simulation model consisted of the empirical movement tracks of nine zebra between 2009 and 2010 (Table 1) and an archive of carcass surveillance records from 1968-2011. The GPS data from the zebra were used to determine the optimal number of distinguishable behavioral states, parameterize the step length and turning angle distributions during each of those states, and calculate the probabilities of transitioning between behavioral states. The carcass surveillance data enabled the initialization of the LIZ layer by informing the likely density of carcasses (0.0135 carcasses/km^2^/year) as well as the expected size and age distributions of locally infectious zones on the landscape (82.7% medium, 14.8% small, and 2.5% large).

### Submodels

*Movement processes:* The movement of agents across the landscape was based upon the behavioral state of the individual at time t. For each possible state, a gamma distribution derived from empirical zebra movement paths was used to generate appropriate step lengths. For behavioral states 1 (resting) and 3 (directed movement), the turning angles were also generated from a circular von Mises distribution derived from the same empirical trajectories. The von Mises was selected because it is a commonly used distribution for simulating or analyzing turning angles in animal movement paths [52, 53, 54, 49]. Movement fixes were made at 20 minute intervals (to match the resolution of the empirical zebra trajectories). Prior to each step, the behavioral state was evaluated and updated based on the probabilities of the observed zebra shifting behavioral states. The relationship between time of day (or more accurately, the time since sunrise, *T*) and current behavioral state was extracted from the empirical movement paths, and these functions were used to shift individuals between behavioral states (transitions not expressed below were not observed in the empirical data, and thus, were treated as impossible in the model):

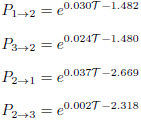

These probabilities served as the basis for Bernoulli draws for each individual in each behavioral state. Under very rare circumstances, the same individual would stochastically be assigned to shift from State 2 to both State 1 and State 3 during the same update. In these cases, the Bernoulli draws were executed repeatedly until the individual was assigned to only one of the two behavioral states.

The foraging state (2) is the behavioral mode most important for the transmission of anthrax. In the simulation model, for individuals in behavioral state 2 at time *t*, movements were made according to the step length distribution observed in empirical movement paths, but the turning angle emerged based on the forage quality in surrounding cells. The perceptual range value (*P_i_*) of individual *i* was used to create a search radius. The individual then adjusted its position to direct its movement toward the center of the cell with the highest relative forage quality (among all of the cells whose center was within the search radius).

*Landscape processes:* Two primary processes affected the landscape, namely feeding and vegetation growth. Feeding occurred each hour (i.e., after three movements by agents), and affected the landscape in two ways. The first form of feeding, called general feeding, occurred at the landscape level (on an hourly basis), where an expected number of “untagged” individuals in the foraging state (approximately 54% of the total ungulate population) were randomly dispersed across cells with a relative forage quality value greater than the mean forage quality (*mu*_*F*_). The density of ungulate agents in this simulation was set at 0.89/km^2^, which was chosen based on the approximate density of medium-sized ungulates in Etosha National Park. Of the proportion of this population that was in the foraging state, 80% were assigned to high quality cells and 20% were assigned to lower quality cells (suggesting imperfect detection of high quality vegetation). Once these individuals were dispersed, feeding decreased the current relative forage quality of the cell (*Q*_*j,t*_) without impacting the carrying capacity of the cell (*K*_*j,t*_), following the equation (valid only for changes sufficiently small to ensure *Q*_*j,t*_ > 0):

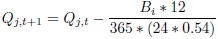

This feeding equation is based on the average biomass consumption of a similar equid species, the horse (*Equus ferus*), per year (3835 - 4146 kg/year; [55]). This value is approximately 12 times the mean body size of the agents in this simulation, so the above equation results in an average daily extraction rate of approximately 11.5 kg/day.

A second, similar form of feeding, called specific feeding, affected only cells that contained a tracked (or “tagged”) individual currently in the foraging state. In those cells, forage quality was also decreased according to the formula above for each individual located therein.

Each hour, a growth process also occurred such that any cell whose current forage quality (*Q*_*j,t*_*S*) was less than the carrying capacity of the cell (*K*_*j*_, as set at initialization) would experience an increase in its quality. The growth process was governed by a logistic equation with a fixed maximum growth rate (*r*, set at 0.10 based on [17]) applied across the landscape:

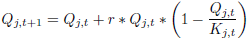

This basic logistic growth equation [56] was used rather than a more complex function (such as that presented in [17, 47]) because the foraging dynamics were not the primary focus of the model. Logistic growth offered a means of maintaining a relatively constant overall landscape quality while offering the ability to implement short-term (i.e., daily) constraints on individual decisions.

## Analysis Methods

The simulation model presented above enables the calculation of probabilities of contact between hosts and LIZs at several different scales of analysis. These analysis methods reveal how alternative scales of movement analysis may contribute significantly, and differently, to our understanding of the epidemiological aspects in question. Though the model does not reflect empirical data directly, the outputs represent a particular construction of the system that can be analyzed in much the same way as field data from a disease system. As such, this agent-based formulation of the anthrax system offers a means of comparing the behavior of alternative methods for estimating important rates that are often difficult to ascertain with empirical data.

### Broad-scale Analysis

The broadest scale of analysis conducted in this case occurred at the level of the home range [57, 58, 59]. This area describes the portion of the park that is utilized by an animal during the course of its normal activities, but this simple definition belies the diversity of methods currently available to demarcate this region. In this case, we have used two different methods: the minimum convex polygon (MCP) [60, 61] and the Local Convex Hull (LoCoH) method [58, 59]. The MCP method is arguably the simplest conception of the home range and depends upon the creation of a polygon that encloses a pre-defined proportion of the points. Though the 95% MCP is frequently used to eliminate outlying points that may be anomalous with regard to the normal activity patterns of an individual, we opted to use the 100% MCP to ensure that contacts with LIZs were not missed. The LoCoH method aims to more accurately reflect the home range by reducing the amount of unused space within the delineated area. Rather than building an MCP around all of the points, the LoCoH algorithm seeks to construct a series of hulls around smaller subsets of points (See Supplementary Materials for details on parameterization of the LoCoH algorithm based on [62]). The result is often more tightly fitted to the known movement track of the individual than the MCP-based home range. Here, too, we used the 100% hullset to verify that all potential points of contact between the animal and the infectious reservoir were considered in our calculation.

**Figure 3:**
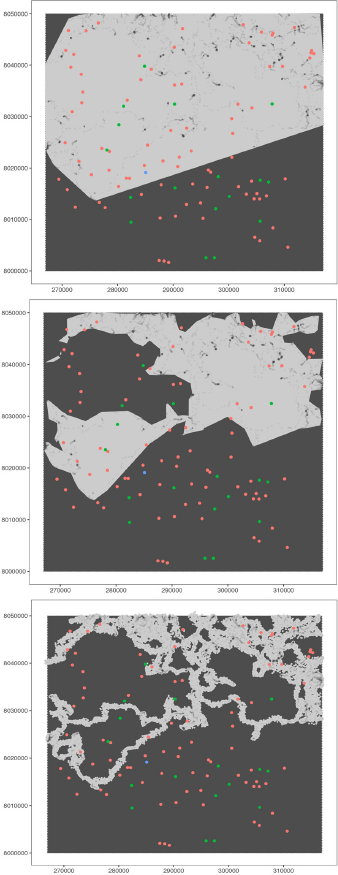
Comparison of the Analysis Methods. The gray regions represent the area considered in the calculation of the probabilities of contact using the MCP (top panel) and LoCoH home ranges (middle panel), as well as the space-time prism (STP; bottom panel) method. The colored points indicate the positions of the LIZs (red dots represent zebra, green springbok, and blue elephants).

### Fine-scale Analysis

A finer-scale analysis of the movement trajectory involved the explicit consideration of uncertainty around each point rather than the more holistic approach of delimiting a home range around the entire path. To address the uncertainty inherent in the discrete positional fixes, we created a set of simulated possible paths to develop a probability surface around the known fix locations. We simulated intermediate points using the space-time prism concept (STP; [63]). This method uses the time budget and the maximum possible speed of movement (*V*_max_) of the individual to simulate a series of steps between the known points that would enable the individual to cover the distance between the origin and destination point in the time allowed. However, rather than a single path in between known fixes, we generated 50 alternative paths [64] of the 19 intermediate points (and one point at each of the known locations). The purpose of this was to enable the creation of a probability surface that was effectively continuous to reflect the continuous nature of movement.

### Considerations of Behavior

Assuming that some knowledge concerning the mode of transmission of anthrax is available, a researcher may conclude that only points obtained when an individual was in the foraging state should be considered in the calculation of the probability of contact. The incorporation of behavioral state analysis may impact the results at both the broad- and fine-scale, so the methods described above were applied to reduced datasets that only included points during which the animal was assessed to most likely be in the foraging state.

To segment our simulated movement trajectories into approximate behavioral states, we used the k-means clustering algorithm of Hartigan & Wong (1979) [65] on individuals’ step sizes seeded with three *a priori* centers (0m, 150m, 1000m) according to visual inspection of individuals’ step size distributions (for example distribution, see Figure 4, right panel). With location data at a resolution of 20 minutes per fix, only canonical activity modes (CAMs; [66]) can be effectively identified, rather than the more specific fundamental movement elements (FMEs; [66]) that underlie these observations. Table 4 reports the mean, median, and standard deviation of step sizes in each cluster, or CAM, averaged across simulated individuals. The k-means clustering algorithm classified 89.1% of the points into the correct behavioral mode, however, it performed even better when only steps in the foraging state were considered, accurately assigning 99.0% of those steps.

**Figure 4:**
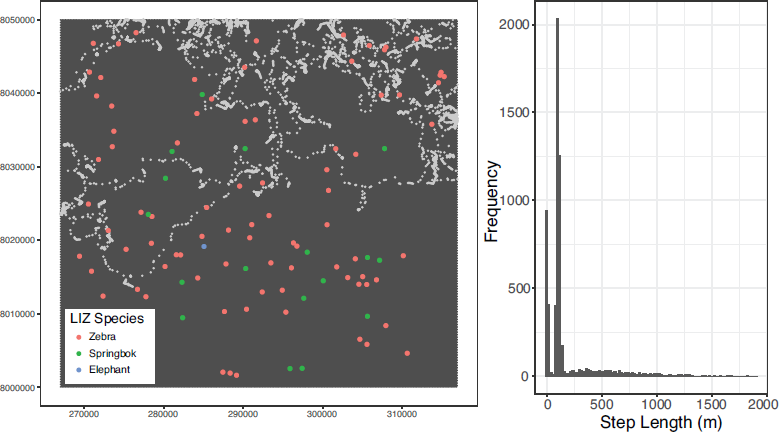
Example Simulated Path. The left panel illustrates the movements of each agent were simulated over the course of 90 days at 20 minute increments for a total of 6481 observations. The trajectory of a single exemplar is plotted here (grey points) as it moves across a landscape, potentially making contact with locally infectious zones (red, blue, and green points) as it moves. The right panel is a histogram of the step lengths of the individual over the entirety of its trajectory.

For subsequent analyses, an animal was considered to be in the foraging state during the 20 minute period between fixes when either the preceding, succeeding, or both fixes were predicted to be in state 2. This method serves as a relatively liberal estimate of time spent in the foraging state (i.e., likely overpredicting foraging points) by including the entire period during which a transition between states was predicted to occur. This, in turn, should offer the most inclusive estimate of potential contact events, such that Type I errors (including a non-foraging time period during which contact occurs; false positive) are more likely than Type II errors (excluding a foraging time period in which contact occurs; false negative).

### Calculation of Contacts

A raster layer of the synthesized locations of LIZs was created with 10 meter by 10 meter grid cells *v*, *v* = 1, …, *w*. The resulting circular buffer polygons were then laid atop the raster, and the proportion of each cell covered by the buffer polygon was calculated to represent the probability of encountering a LIZ in each cell. To calculate the probability of contact for each individual, this synthesized LIZ layer was considered in conjunction with the probability surfaces created by the broad- and fine-scale analyses. The surfaces obtained after the explicit incorporation of behavioral state were also considered to determine the utility of the additional analysis step at both scales. Because of the uncertainty regarding the dose required to cause infection or death in a wild ungulate, we considered only the probability of a transmission-enabling contact rather than the actual probability of transmission itself.

In the case of the broad-scale analyses (with and without behavior considered), the home ranges that emerged from the MCP and LoCoH methods were first rasterized onto the same 10 meter by 10 meter resolution grid, and then set down upon the LIZ layer. Because the entire home range was treated as having a uniform probability of presence throughout, the number of cells encapsulated by the home range was calculated (including the portions of cells covered at the edges) and the probability of 1 was distributed accordingly. If 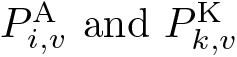 represent the respective probabilities of an agent foraging in cell v and of a carcass being in cell *v*, then the probability of individual *i* encountering at least one of the z LIZs during the anthrax season 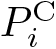 was calculated as:

**Table 3:** Basic descriptive statistics of step lengths for simulated individuals during each of the potential behavioral states.

| State | Median | Mean | StDev |
| --- | --- | --- | --- |
| Resting | 6.38 | 8.57 | 7.76 |
| Foraging | 105.7 | 113.5 | 100.0 |
| Traveling | 545.1 | 630.5 | 404.9 |

**Table 4:** Basic descriptive statistics of step lengths for empirical individuals during each of the behavioral states defined by the three-state hidden Markov model.

| State | Median | Mean | StDev |
| --- | --- | --- | --- |
| Resting | 5.60 | 7.96 | 7.46 |
| Foraging | 68.4 | 104.2 | 111.9 |
| Traveling | 577.5 | 647.1 | 398.8 |

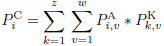

For the fine-scale analyses, the simulated points were summarized using another 10 meter by 10 meter resolution raster. The number of simulated points within each raster cell was calculated and divided by the total number of simulated points across the entire path (6,805,050 points in the case of the full path of an individual). By dividing by the total number of points, the sum of the values of all of the raster cells was equal to 1 for each individual, thereby representing a probability of presence over the course of the anthrax season. The probability surface derived from the simulated paths was used instead of the home range raster, but the same equation was applied.

## Results

LIZs occupied at least some portion of 422 of the raster cells (10 by 10 meters) on the landscape. That represents a proportion of 0.000014 of the total number of cells in the park. The probabilities of contact for every individual, irrespective of analysis method, are greater than that expected through random movement alone (Figure 5; Table 5). The probability of contact values presented below effectively represent the proportion of steps during an anthrax season that an individual will be present in a cell with anthrax spores.

**Figure 5:**
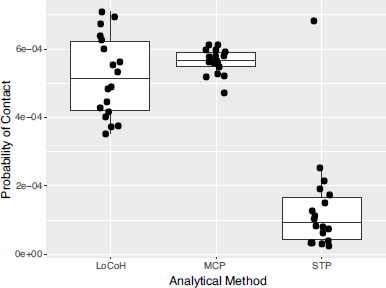
Probability of Contact Estimates Across Analysis Methods. A boxplot of the probabilities of contact that emerged from the three alternative analysis methods: LoCoH (leftmost box), MCP (middle box), and STP (rightmost box). The black dots indicate the probabilities of contact associated with the 18 simulated agents, though their dispersion along the x-axis is merely for clarity within the three analysis types.

The MCP-based broad scale analysis method applied to the full dataset resulted in estimates of probabilities of contact that ranged from 0.000471 to 0.000613 (mean = 0.000564; SD = 0.000037). Estimates obtained using the reduced dataset, representing only those points during which the individual was in the foraging behavioral state, were almost identical (mean = 0.000563; SD = 0.000037), with only two individuals estimated to have different probabilities of contact (IDs 9 and 14).

The LoCoH-based broad scale analysis resulted in substantially different estimates. Using the full dataset, the probabilities of contact ranged from 0.000353 to 0.000709 (mean = 0.000521; SD = 0.000118). This level of heterogeneity between individuals was not reflected in the more simplistic MCP approach. In addition, the application of the LoCoH-based method to the foraging-only dataset resulted in significantly lower estimates of probabilities of contact ( *p* = 0.0012 based on a paired t-test). The values obtained using the reduced dataset ranged from 0.000175 to 0.000693 (mean = 0.000402; SD = 0.000132). On average, the probabilities of contact using the reduced dataset were 21.6% lower than those estimated from the full dataset.

The finer-scale analysis based on the space-time prism approach resulted in estimates of probabilities of contact ranging from 0.000026 to 0.000682 (mean = 0.000137; SD = 0.000152). The values obtained using the reduced dataset ranged from 0.000015 to 0.000366 (mean = 0.000091; SD = 0.000087). Like the LoCoH-based analysis, the differences between the probabilities derived from the full and reduced datasets were significantly different (*p* = 0.03 based on a paired t-test). On average, the probabilities of contact using the reduced dataset were 28.5% lower than those estimated from the full dataset.

The individual-based methodology resulted in estimates that were significantly lower than either home-range-based approach. Paired t-tests between the estimated probabilities of contact based on the STP method relative to the MCP method demonstrate significant differences for both the full dataset (t = 10.93; *p* < 0.0001) and the reduced datasets (t = 18.9; *p* < 0.0001). Comparing the results from the LoCoH approach to the STP method reveals similar patterns for both the full (t = 7.64; *p* < 0.0001) and reduced (t = 8.05; *p* < 0.0001) datasets. Interestingly, though the probabilities of contact estimates from the MCP method do not differ significantly from the LoCoH method when the full dataset is considered (t = 1.64; *p* = 0.12), the differences are significant when the behavioral component is considered (t = 5.32; *p* < 0.0001). This suggests that the inability of the MCP to respond to the removal of epidemiologically-irrelevant points could lead to dramatic overestimates of the probability of contact during susceptible behavioral states.

**Table 5:** Probability of contact values across 18 simulated agents using three alternative estimation methods: Minimum Convex Polygon (MCP), Local Convex Hull (LoCoH), and Space-time Prism (STP), each applied to the full trajectory (Full) and a subset of the dataset during which the individual was predicted to be in the foraging behavioral state (Forage).

| Simulant ID | MCP |  | LoCoH |  | STP |  |
| --- | --- | --- | --- | --- | --- | --- |
|  | Full | Forage | Full | Forage | Full | Forage |
| ID1 | 559* | 559 | 353 | 175 | 35 | 18 |
| ID2 | 593 | 593 | 628 | 486 | 80 | 79 |
| ID3 | 599 | 599 | 696 | 257 | 62 | 054 |
| ID4 | 612 | 612 | 554 | 624 | 173 | 122 |
| ID5 | 558 | 558 | 483 | 289 | 32 | 24 |
| ID6 | 471 | 471 | 446 | 344 | 114 | 83 |
| ID7 | 563 | 563 | 601 | 416 | 103 | 101 |
| ID8 | 548 | 548 | 430 | 406 | 26 | 15 |
| ID9 | 568 | 554 | 401 | 326 | 150 | 150 |
| ID10 | 599 | 599 | 491 | 290 | 76 | 67 |
| ID11 | 613 | 613 | 416 | 359 | 39 | 24 |
| ID12 | 579 | 579 | 377 | 467 | 191 | 150 |
| ID13 | 523 | 523 | 563 | 321 | 215 | 200 |
| ID14 | 528 | 522 | 374 | 341 | 682 | 366 |
| ID15 | 519 | 519 | 535 | 438 | 126 | 92 |
| ID16 | 567 | 567 | 639 | 693 | 82 | 26 |
| ID17 | 580 | 580 | 675 | 431 | 254 | 36 |
| ID18 | 580 | 580 | 709 | 573 | 33 | 27 |

\* All values in this table must be multiplied by $10^{-6}$ : i.e. this first entry is 0.000559

## Discussion

To demonstrate the benefits of incorporating movement data into disease research, we analyzed the outputs of an agent-based simulation model. The goal of our exercise was to estimate, using some of the tools described above, the probability of individuals coming into contact with an infectious dose of an indirectly-transmitted pathogen within an environmental reservoir. The outputs of the simulation included a set of movement trajectories (covering the three-month anthrax period at a temporal resolution of 20 minutes per fix) and the locations and sizes of LIZs across the landscape. Due to the limitations of most field surveillance efforts, it is unlikely that a researcher would have the ability to map out the exact locations of every infected carcass on the landscape (estimates from the Etosha system based on a hierarchical model of distance sampling place the rate of detection at approximately 25%; [45]), so the simulation framework offered an alternative approach that enabled full knowledge of the distribution of risk across the landscape. Thus a comprehensive map of LIZ sites was used for the estimation of contact rates to judge the relative strengths of various movement analyses at different scales.

The analyses conducted here demonstrate the value of using fine-scaled movement data in estimating rates of epidemiological relevance. The use of methods that function at the scale of the home range resulted in higher probabilities of contact than methods that adhered more closely to the movement trajectory of the individual. The general purpose of most home range delineation methods is to generalize from relatively sparse movement paths, so the inclusion of large areas is not surprising, nor are the correspondingly high estimates of contact rates. The space-time prism method, on the other hand, enabled the incorporation of some level of uncertainty regarding the movement of the animal in between GPS fixes without generalizing too far beyond the measured path. Thus, the area used for calculating the probability of contact was less likely to include a large number of LIZs. Even using the simulation framework where the locations of LIZs are known, there remains no means of extracting a ‘true’ probability of contact for each individual. Rather, we can demonstrate the performance of these alternative analysis methods in a controlled microcosm that would be difficult to replicate in the field.

In most wildlife disease systems, precise contact rates, whether generalized over an entire host population or recorded for each individual separately, are only very rarely calculated directly. The effort involved in the near-continuous monitoring of individuals normally precludes such calculations (but see fairly extensive studies regarding tuberculosis transmission among badgers and livestock; e.g., [67, 68]). Here we demonstrate that relatively straightforward movement analyses can be used as a proxy for intensive sampling regimes, offering insight into the magnitude of variance among individuals as well as average rates for use in more traditional epidemiological models. Analyses applied at the home range scale tended to result in larger estimates of contact rates than those applied at the individual path level. When considering the ultimate impact on the transmission term *beta* frequently fitted to epidemic data, any overestimation of contact results in a reciprocal underestimation of the infectiousness of a pathogen. Similarly, underestimation of the variance among individuals, as results from the MCP-based approach, could lead to incorrect conclusions about the variance in potential outbreak sizes. For example, the rapid fade-out of an epidemic may be attributed to the pathogens infectiousness (or lack thereof) when, in fact, it is merely the result of a stochastic movement process. The same heterogeneity that led to fade-out after one introduction of infection could very well give rise to a much larger epidemic following another introduction into the same population.

Due to the probabilistic nature of the analyses at the individual and landscape levels, all 18 of the simulants were estimated to have had some non-zero probability of encountering a LIZ. Field-based investigations of sub-lethal exposure in ungulates in Etosha revealed that 52%-87% of sampled zebra exhibited some level of anti-anthrax antibody titres [38], though the rate at which titres may wane over time is not known. Nevertheless, while the analysis-derived values may be slight overestimates, it is not unreasonable to expect that every individual may come into contact with at least a few anthrax spores, perhaps following deposition by tabanids (horse-flies) on browse [41]. Differences in the infectious doses encountered by each individual—the cores of LIZs versus a few stray spores— may be one possible explanation for the majority of individuals exhibiting some level of exposure.

The LIZs in our simulation were relatively small, with infectious radii ranging from about 1 meter (for the smallest carcasses, akin to a springbok) to about 14 meters (for the rare large carcasses, akin to elephants). These infectious sites occupied a total area of approximately 13,000 m^2^, or about 0.013 km^2^. Even so, all 18 of the simulated individuals were predicted to encounter at least one LIZ along their movement trajectories during the anthrax season. This is likely because of the attractive nature of these areas, which increase the forage quality in the vicinity of the carcass [69, 34], a feature that emerges from the movement dynamics during the foraging phase of the simulation.

Several of the complexities of infection dynamics, including considerations of heterogeneity in dosages and immune responses, were excluded from this model. Instead, the emphasis was on potential transmission events and a more readily measurable metric: the probability of contact with a LIZ bounded by the radii mentioned above. For this reason, the relatively short-time frame of the simulation was selected. Though the peak anthrax season is likely long enough for some individuals who encounter infectious carcass sites to succumb to the disease, such mortality events were not incorporated. This was deemed reasonable, as such newly deposited carcasses have been reported to have repulsive effects on live individuals [34], meaning that they would be unlikely to influence the subsequent probabilities of contact for monitored agents. Several additional state variables and submodels would be needed to consider more complete infection dynamics. Additional state variables would include parameters defining agent immune systems and changing bacterial densities at LIZs. Infection and immune response submodels would also be required, which could account for altered movement patterns in infected hosts and disease-induced mortality during and after the anthrax season. Due to the lack of empirical data for parameterizing such state variables and submodels, the model presented here was not extended in this manner.

According to the pattern-oriented modeling framework [13], multiple observed patterns should be matched by an agent-based model to optimize model structure, compare alternative rule sets, and reduce parameter uncertainty. The fact that infection with anthrax is difficult to ascertain prior to an animal succumbing means that there are not many empirical metrics to use for verification of the model outputs. To verify that emergent properties arise from the model, a more theoretical pattern was matched: changes in the density of foraging agents on the landscape should alter the probabilities of contact in a directional, but potentially non-linear, fashion. This is based on the idea that additional agents (i.e., increased densities) would be similarly attracted to LIZ sites, extract resources from those areas, and reduce the likelihood of tagged agents foraging at such sites, thereby decreasing the mean probability of contact for all tracked individuals. This theoretical pattern was tested with the simulation model at three densities of hosts: *D* = 0.495, 0.89, and 1.78 agents/km^2^ (representing the predicted density as well as one-half and twice that density). Though the relationship is not statistically significant at the 0.05 level (*p* = 0.10), there exists a downward trend in contact probability with increased agent density that may be borne out with greater clarity under a wider range of densities (see Supplementary Figure S2).

Ultimately, the model serves as a proof of concept that the incorporation of movement data collected using available technology may help elucidate some of the contributing aspects of infection processes. For pathogens with an environmental reservoir, like *B. anthracis*, evaluating the probability of susceptible hosts contacting sources of infectious material is likely to be one of the most valuable means of assessing the risk faced by local wildlife populations. Real world pathogens with dependence on readily remotely-sensed environmental factors may be more amenable to analyses at broader scales, as sites of pathogen deposition may be readily predicted from imperfect or incomplete data. In such cases, the necessary resolution of the movement data may be relaxed while maintaining some ability to draw conclusions about individual risk of infection.

Despite the data-intensive nature of agent-based models, they hold a great deal of promise for understanding the complex dynamics underlying epidemics, particularly in wildlife populations. The use of empirical movement data to parameterize this simulation model also demonstrates the value of incorporating such increasingly ubiquitous data sources. These methods may help guide future data collection efforts or elucidate certain traits (e.g., habitat preferences) that indicate heterogeneous vulnerability among hosts. Thus, such combinations of tools can alter the means by which risk is evaluated in disease systems, such as those dominated by anthrax.

## Acknowledgements

The authors would like to thank the Getz and Brashares labs at UC Berkeley for their helpful comments. This research was supported by grants NIH GM083863 (WMG) and NIH GM117617 (PI Jason Blackburn; ED and DS supported under a UC Berkeley subcontract).

